# Early and late evoked brain responses differentially reflect feature encoding and perception in the flash-lag illusion

**DOI:** 10.1101/2021.06.03.446877

**Authors:** Julian Keil, Daniel Senkowski, James K. Moran

## Abstract

In the flash-lag illusion (FLI), the position of a flash presented ahead of a moving bar is mislocalized, so the flash appears to lag the bar. Currently it is not clear whether this effect is due to early perceptual-related neural processes such as motion extrapolation or reentrant processing, or due to later feedback processing relating to postdiction, i.e. retroactively altered perception. We presented 17 participants with the FLI paradigm while recording EEG. A central flash occurred either 51ms (“early”) or 16ms (“late”) before the bar moving from left to right reached the screen center. Participants judged whether the flash appeared to the right (“no flash lag illusion”) or to the left (“flash-lag illusion”) of the bar. Using single-trial linear modelling, we examined the influence of timing (“early” vs. “late”) and perception (“illusion” vs. “no illusion”) on flash-evoked brain responses, and estimated the cortical sources underlying the FLI. Perception of the FLI was associated with a late window (368-452ms) in the ERP, with larger deflections for illusion than no illusion trials, localized to the left fusiform gyrus. An earlier frontal and occipital component (200-276ms) differentiated time-locked early vs. late stimulus presentation. Our results suggest a postdiction-related reconstruction of ambiguous sensory stimulation involving late processes in the occipito-temporal cortex, previously associated with temporal integration phenomena. This indicates that perception of the FLI relies on an interplay between ongoing stimulus encoding of the moving bar and feedback processing of the flash, which takes place at later integration stages.

**Highlights:** Flash-lag illusion relates primarily to late evoked brain potentials (>300ms)

Illusion vs. no-illusion trials showed difference in fusiform gyrus

Flash-lag illusion could involve postdiction-driven integration of ongoing stimuli

## 1 Introduction

In our environment, incoming stimuli are continuously integrated into the ongoing stream of sensory inputs, leading to smooth conscious perception. Some perceptual illusions could provide a clue as to how this is performed. In the case of the Flash-Lag illusion (FLI), when a moving object is presented with a briefly flashed stimulus, the moving object is misperceived as being further along its trajectory than it really is (Nijhawan, 1994). Although there are a number of theoretical assumptions regarding the origins of the FLI, many have in common the idea of a designated time window, within which the moving object and the flash could be integrated (Hubbard, 2014). Perception is shaped by input preceding the flash stimulus and there is likely also a time window of a few hundreds of milliseconds within which information that is presented after a stimulus can retroactively affect the perception of this stimulus (Sergent, 2018; Shimojo, 2014). Theories differ in terms of the relative weighting of the pre-flash and post-flash sensory processing for the FLI. In a parallel manner, the proposed neural mechanisms underlying these theories are also separate, with some emphasizing early, temporally stimulus-locked processing (e.g. Hogendoorn, 2020) and others emphasizing later stimulus decoupled global processing (Sergent, 2018).

Theories focusing on the importance of pre-stimulus sensory processing for perception have linked the FLI to interactions between the higher visual area (V5) and the primary visual cortex (V1). One of the most thoroughly elaborated theories, both in theoretical and empirical terms, is the motion extrapolation theory (Hogendoorn, 2020), in which the window of integration is reflected in higher level visual areas, which are preactivated in anticipation of ongoing movement, as a means of compensating for temporal lags in neural transmission from lower- to higher-level feedforward connections (Hogendoorn & Burkitt, 2019). In these terms there is a disjunct between the anticipated movement of the moving stimulus and the unpredictable onset of the flash stimulus. Recently, Hogendoorn & Burkitt, (2018) contrasted predictable vs. non-predictable moving stimuli and found that the former could be decoded from relatively early EEG activity, i.e., around 140ms after stimulus onset. Moreover, a functional neuroimaging study showed that motion-related stimulus processing in V5 and V1 appears to be subject to predictive coding, with less predictable visual movements producing greater BOLD responses (Schellekens et al., 2016). Another elegant explanation of the FLI is the non-linear latency difference theory (Arstila, 2015), which involves reentrant processing from V5 to V1. In this theory, reentrant processing from V5 to V1 is related to conscious perception. The FLI is predicted to stem from a violation of this process: the stimuli create a conflict between the reentrant processing of the early stimulus, i.e., the moving bar, and feedforward processing of the later flash stimulus in V1, leading to the illusory perception of a lagging flash. In sum, there are compelling theories linking FLI to early (<200ms) visual prestimulus processing, tightly coupled to the order of stimulus presentation.

In contrast to these theories, the concept of postdiction emphasizes the importance of information that follows the flash. Perception phenomena in which a second stimulus retroactively affects the perception of a first stimulus have been coined postdiction (Shimojo, 2014). Foundational findings in support of the postdiction hypothesis come from Eagleman & Sejnowski, (2000), who posit that the window of integration is biased by the subsequent motion of the stimulus. In support of this, the authors demonstrated that the FLI is maintained and can even be further manipulated when only a post-flash movement is present, but far less so when only pre-flash movement is present (Eagleman & Sejnowski, 2007). Postdiction is currently operationally defined, and agnostic as to the underlying neural mechanisms. However, integration across greater timescales, decoupled from the actual temporal order of stimulus presentation, will likely require an involvement of top-down processes (Sergent, 2018). Thus, in contrast to theories of early stimulus-locked neural processing, postdiction is more compatible with global top-down processing at later integration stages.

Other temporally governed perceptual phenomena show a split between early and later components in the empirical literature (Förster et al., 2020). They have been invoked to support competing theories. For example, backward-masking shows the dependency of conscious perception on reentrant processing between V5 and V1 (Fahrenfort et al., 2007). However, Cul et al., (2007) found that the difference in backward-masked trials subjectively rated as invisible vs. visible, was not related to early P1 or N1 components, but rather to the later P3 component. In a parallel manner, retroactively altered perception of basic stimuli in the FLI may be dependent on either earlier local-recurrent loops or on later top-down reamplification across wider areas of the brain.

A more precise picture of the time course of cortical activation would help clarify the relative strengths of the above theoretical accounts on perception in the FLI. Up until now, however, there has been little investigation of the phenomenon using methods with a fine time resolution, such as EEG or MEG. One EEG study from Stekelenburg & Vroomen, (2005) has shown the effects of an audiovisual manipulation of the visual flash lag, which reduced the flash-lag effect and a correspondingly reduced N1 event-related potential (ERP) component. Moreover, the FLI can be disrupted at approximately 200ms post flash by TMS stimulation of MT+ (Maus et al., 2013). In another EEG study, Chakravarthi & VanRullen (2012) examined the single-trial oscillatory correlates of FLI, showing that the illusion was dependent on the phase of alpha and beta oscillations pre- and post-flash onset, respectively. This suggests the periodic sampling of temporal windows rather than the continuous sampling of individual time points. Still, an ERP study comparing flash-lag vs. non-flash-lag processes directly across a longer time scale is still missing from this literature.

In the present study, we examined the ERP components of a FLI, testing directly the difference in the ERPs between stimuli where the flash-lag is perceived and where it is not perceived. Differences in early components (C50, P1, N1 <200ms) would be congruent with theories emphasizing early processing, e.g., motion extrapolation or reentrant processing, whereas differences in later components (>300ms) would support theories that emphasize later feedback processing of postdiction.

## 2 Methods

### 2.1 Participants

An initial sample of 29 paid volunteers participated in this study. All had normal or corrected to normal vision and reported no history of neurological or psychiatric illness. Twelve participants were removed from the sample after participation: Six were excluded due to noisy EEG data or insufficient detection of the catch trials, and six were excluded because they had too few trials in the examined conditions (e.g., too few or too many illusion trials, see below). After preprocessing of the electrophysiological data, 17 participants (8 female, mean age: 38.47 (Range: 24-52)) were included in the final data analysis. Handedness 15 were right-handed according to the Edinburgh Handedness Inventory, (Oldfield 1971), with two <50% right-handed. The study was conducted in accordance with the local Ethics Committee of the Charité – Universitätsmedizin Berlin as well as with the 2008 Declaration of Helsinki, and all participants provided written informed consent (Ethical approval number: EA1/169/11).

### 2.2 Task

Participants were seated in a dimly lit, electrically and acoustically shielded chamber, while being presented with stimuli of the flash-lag paradigm. Participants had to indicate by a button-press whether a visual flash appeared to the right or to the left of a moving bar (Fig. 1A). On a CRT monitor, a black bar (1.33° x 0.28° visual angle) moved from the left to the right with a speed of 10° visual angle per second for 1400 ms. The distance from the bar onset to the screen center was 6.74° visual angle, and the bar reached the screen center after 700 ms. A fixation cross was presented for the whole trial at the bottom center of the screen. Above the fixation cross (3.05° visual angle), a white circle (0.28° visual angle) flashed for 16.7 ms, either if the bar was 51 ms (*early* trials) or 16 ms (*late* trials) away from reaching the center of the screen. After the bar reached the right side of the screen, the bar remained stationary for 100 ms before disappearing and the fixation cross turned into a hand symbol as a response cue. The response cue ensured that there was no confounding motor activity during the stimulation period. The response interval had a random duration between 1500 ms and 2500 ms. Additional audiovisual trials were presented, which contained a 16.7 ms 72 dB(SPL) white noise burst 100 ms before the bar reached the screen center (Stekelenburg and Vroomen, 2005). These trials were not entered into the data analysis because they were not relevant for the current research question. Overall, the experiment consisted of 200 trials per condition (early flash without a burst; late flash without a burst; early or late flash with noise burst), 168 trials (equally distributed across conditions; ~15% of all presented trials) with a reversed direction (right to left) of the bar movement, and 172 catch trials (equally distributed across conditions; ~15% of all presented trials) in which the central fixation cross was surrounded by a box with the cross within it. This change in catch trials occurred 60 ms before the bar reached the center of the screen and participants were required to respond to the trials by pressing both buttons simultaneously. Trials with a reversed movement were included to avoid habituation effects. Catch trials were included to ensure that participants focused on the central fixation cross. Trials of the different experimental conditions, reversed direction trials, and catch trials were presented in random order. In sum, 1140 trials were presented. Each trial had a duration of 3500 ms, leading to a total experimental runtime of about 66 minutes (divided into 15 blocks).

**Figure 1:**
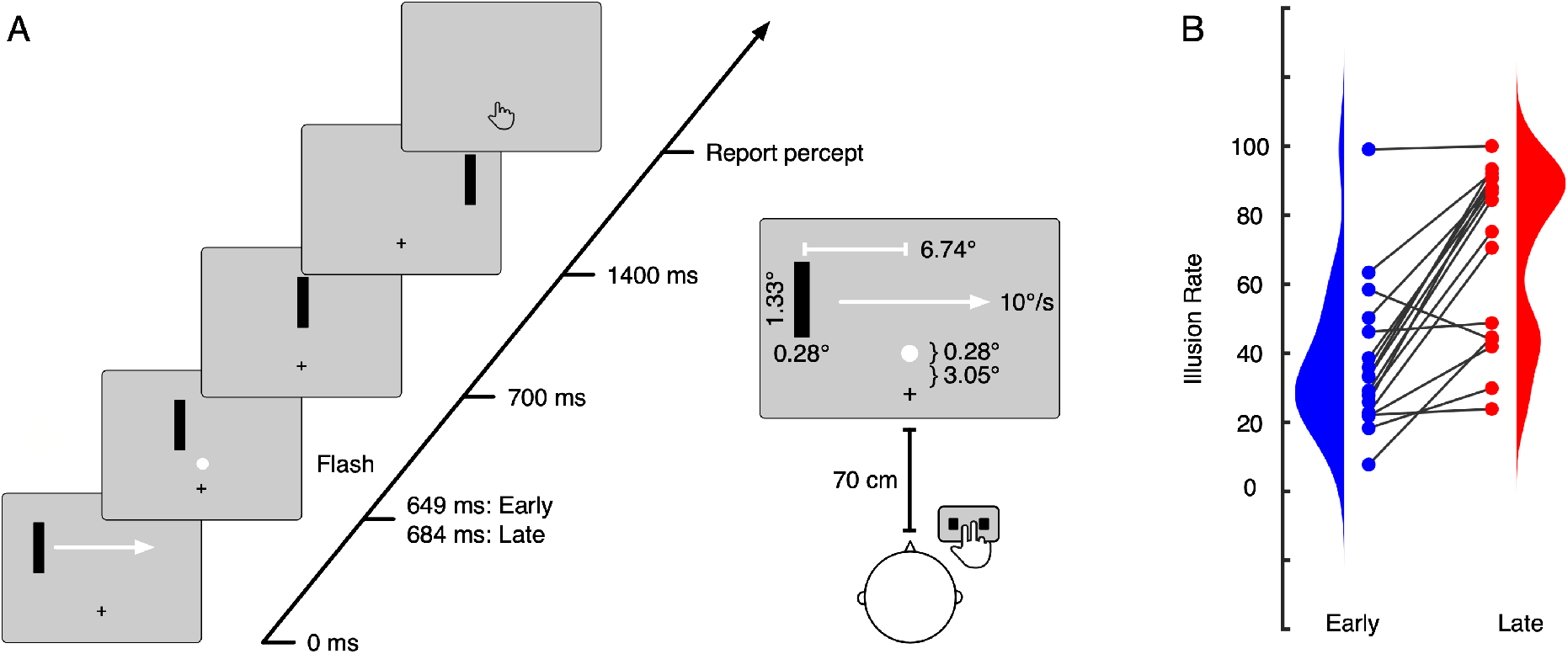
Experimental setup of visual flash-lag trials and behavioral findings. (A) A black bar moved from left to right across the screen for 1400 ms. Either 51 or 16 ms before the bar reached the center of the screen, a white circle was flashed for 16.7 ms. Participants had 1500-2500 ms to respond using the right hand. (B) Participants perceived the flash-lag illusion more often when the flash appeared late (i.e., 16.7 ms prior to the bar reaching the center; red) compared to when the flash appeared early (i.e., 51 ms prior to the bar reaching the center; blue).

### 2.3 Behavioral Data Analysis

Participants reported their perception with their right hand using a CEDRUS response pad, with a left button pressed by the index finger as the circle appearing ‘left’ relative to the bar and a right button pressed by the middle finger as the circle appearing ‘right’ relative to the bar and both buttons simultaneously as soon as the catch trial cue appeared. From the participants’ responses, trials were categorized as *illusion trials* if participants reported the central flash on the left of the bar moving from the left to the right of the screen, i.e., if the central flash perceptually lagged the moving bar and as *no illusion trials,* if they correctly reported the flash to be to the right (relative to the moving bar) when the flash was presented.

### 2.4 EEG Data Analysis

EEG data were recorded with a high-density 128-channel EEG system (EasyCap, Herrsching, Germany), including one horizontal and one vertical EOG electrode placed below and lateral to the right ocular orbit to register eye movements using Brainamp DC amplifiers (Brainproducts, Gilching, Germany). Recordings were made against nose reference at a sampling frequency of 1000 Hz and with a passband of 0.016–250 Hz.

EEG data preprocessing and data analysis were conducted in MATLAB (MathWorks, Natick, MA, USA) using EEGLAB (http://www.sccn.ucsd.edu/eeglab) (Delorme & Makeig, 2004), FieldTrip (http://www.ru.nl/fcdonders/fieldtrip) (Oostenveld, Fries, Maris, & Schoffelen, 2010) and customized scripts. First, data were filtered offline using windowed sinc FIR filters (Widmann et al., 2015) (high pass: 1 Hz, low pass: 125 Hz, notch: 49–51 Hz). Furthermore, data were down sampled to 500 Hz and epochs from − 1 to 1 s relative to flash onset were extracted from the data. Epochs with large artefacts were removed by visual inspection (M removed trials = 222.00, SD = 169.90). To further correct for EOG artefacts (blinks, muscle activity) and strong cardiac activity, independent component analysis (ICA) was conducted (Runica, Lee et al., 1999). On average 3.65 ICA components (SD = 1.32) were removed. Channels with extremely high artifacts were interpolated with distance interpolation (M removed electrodes = 7.65 electrodes, SD = 3.41). The EOG channels were not included in the further analysis of ERPs. For the analysis of ERPs, epochs were lowpass filtered below 45 Hz using windowed sinc FIR filters (Widmann et al., 2015), and the epoch mean was subtracted. For the EEG data analysis only trials in which the bar moved from the left to the right were used. After artifact rejection on average 81.4 (SD +/− 35.27) trials were available (*early illusion* = 119.7 +/− 44.51, *early no illusion* = 47.0 +/− 36.85, *late illusion* = 56.29 +/− 29.83, *late no illusion* = 102.64 +/− 37.23).

To investigate the cortical sources of the observed ERP responses in the electrode-level analysis, we followed previously established analysis pipelines (Keil et al., 2017; Speer et al., 2020) and performed source localization using a linearly constrained minimum variance (LCMV) beamformer algorithm (Van Veen et al., 1997). A leadfield was generated using a realistic three-shell boundary-element volume conduction model based on the MNI standard brain (MNI; http://www.mni.mcgill.ca) for each grid point in the brain on a regular 10-mm grid. Within each participant, we first constructed a common spatial filter across all conditions from the covariance matrix of the averaged single trials at electrode level and the respective leadfield. The use of a common spatial filter for all data guaranteed that differences in source space activity could be ascribed to power differences in the different conditions and not to differences between filters. The lambda regularization parameter was set to 10%, to compensate for potential rank reduction during preprocessing. We then projected the single condition ERPs for the two time-windows identified in the electrode-level analysis into source space using the precomputed common filter. A baseline correction was performed for each time window and condition, using the inverse interval prior to stimulus onset for each time window. To this end, the activity in the respective baseline interval was first subtracted from the post-stimulus interval of interest, and the resulting difference was then divided by the average baseline activity. The anatomical regions of the source localization were determined based on the automated anatomic labelling atlas (AAL, Tzourio-Mazoyer et al., 2002).

### 2.5 Statistical Analyses of EEG Data

The influence of the flash onset latency and the perception of the illusion on the event-related potentials was simultaneously evaluated using single-trial linear models in the first 500 ms after flash onset. In the first level, the single trial amplitude of the ERP was related to the within-subject factors Time (*early* vs. *late*), Illusion (*illusion* vs. *no illusion*), and the interaction between both factors in each participant. In the second level, beta values for the two factors and the interaction were compared to zero across participants. To this end, we conducted a non-parametric cluster-based permutation test that addresses the multiple comparison problem by clustering together samples adjacent in time and space (Maris & Oostenveld, 2007). The experimental cluster-based test statistic was evaluated against a permutation distribution in order to test the null hypothesis of no difference between beta values and zero using a two-tailed dependent-samples test. The critical alpha level was set to 0.05. In order to control for multiple comparisons across the 2-dimensional matrix of 126 (electrodes)*251 (samples) comparisons, the clustering algorithm searches for neighboring elements below the critical alpha level and sums the t-values in these clusters. Then the condition labels are shuffled, and the same comparison is computed on the shuffled data. This shuffling step is repeated for 1000 iterations, and for each iteration the largest sum of t-values is retained (‘maxsum’ setting). Finally, the t-value sums in the clusters of the empirical data are compared to the distribution of the clusters obtained in iterations. The p-value for each empirical cluster thus is a percentile indicating the likelihood to obtain a cluster of this size based on randomly shuffled data. Importantly, the clusters obtained in this analysis are not due to any á priori selection of a time interval but are the solely based on the empirical data. In order to further compare the ERP between the different factors, trials were averaged within the clusters identified in the previous steps within each participant. Then, we compared ERP amplitudes between *early* vs. *late* and *illusion* vs. *no illusion* trials using parametric paired-sample t-tests. Additionally, we correlated the illusion rate with the ERP amplitude using a non-parametric Spearman correlation. Moreover, to explore the possibility of further ERP differences between conditions, we computed a 2×2 repeated-measures ANOVA for the two-dimensional channel by time space with FDR correction for multiple comparisons. The alpha-level was set to 0.05 in all post-hoc exploratory analyses. Bayes Factors were computed (BF10, Rouder et al., 2009) as an indicator of the relative evidence for the H0 and H1 on the power averaged within the clusters identified in the previous steps. A BF10 between 1 and 3 indicates anecdotal support for the alternative hypothesis (H1), whereas a BF10 between 3 and 10, and above 10 indicate respectively moderate and strong support for H1. A BF10 of 1 indicates equal support for H1 and the null hypothesis (H0) while, on the other side, a BF10 between 1/3 and 1, 1/10 and 1/3 and below 1/10, provides respectively anecdotal, moderate and strong support for H0 (Aczel et al., 2017).

In source space, we again used the aforementioned non-parametric cluster-based permutation test to compare source space activity for the factors Time (*early* vs. *late*) and Illusion (*illusion* vs. *no illusion*) based on 10000 permutations. Source space activity averaged across identified clusters was again correlated with the illusion rate using a non-parametric Spearman correlation. The alpha-level was again set to 0.05 in all post-hoc exploratory analyses in source space, and Bayes Factors were computed as an indicator of the relative evidence for the H0 and H1.

## 3 Results

### 3.1 Behavior

The likelihood to perceive the FLI was influenced by the temporal distance between the bar and the flash (t(16) = −5.5091, CI = [−0.4551 −0.2022], p < 0.001, BF10 = 548.0798). Replicating the findings previous studies (Stekelenburg & Vroomen, 2005), the occurrence of the early flash (Figure 1B blue, 37.28 +/− 21.55, mean % +/− SD) resulted in less illusions than the late flash (Figure 1B red 70.14 +/− 25.30). Participants correctly identified the catch trials (87.04 +/− 16.88).

### 3.2 ERPs

The influence of the flash onset latency and the perception of the illusion on the evoked brain potentials was evaluated using linear modelling within each participant. Then the beta values for the factors Time (*early* vs. *late*), Illusion (*illusion* vs. *no illusion*), and the interaction between the two factors were statistically compared to zero across participants.

For the factor Time, the cluster-based permutation test revealed an interval between 200 and 276 ms during which the beta values differed from zero. This suggests that in this interval, the single-trial ERPs differ between *early* and *late* trials. Specifically, a cluster of negative beta values was found across frontal electrodes (cluster-p = 0.005 +/− 0.0044), and a cluster of positive beta values was found across occipital electrodes (cluster-p = 0.011 +/− 0.0065). Averaging the ERP amplitudes across the electrodes of the negative cluster, trials in which the flash occurred early (Figure 2A blue, 0.9257 +/− 0.5831, mean μV +/− SD) compared to trials in which the flash occurred late (Figure 2A red, 0.9310 +/− 0.6350) were associated with numerically reduced positive ERP amplitudes. However, this difference was not statistically significant (t(16) = −0.0991, CI = [−0.1183 0.1077], p = 0.9223, BF10 = 0.2500). Averaged across the positive cluster, trials in which the flash occurred early (Figure 2E blue, −0.7983 +/− 0.5706) were associated with a numerically less negative ERP amplitudes than trials in which the flash occurred late (Figure 2E red, −0.8149 +/− 0.5738). This difference was not significant (t(16) = 0.3063, CI = [−0.0983 1.1315], p = 0.7633, BF10 = 0.2594). Finally, the correlation analysis between the ERP amplitude averaged over the respective clusters and the illusion rate was not significant (occipital cluster: r(15) = 0.3971, p = 0.1156, BF10 = 0.6361; frontal cluster: r(15) = −0.3260, p = 0.2014, BF10 = 0.4146). Taken together, comparing the beta values obtained from the linear modeling of single-trial ERP amplitudes to zero suggests that ERPs differ between early and late trials. However, the less sensitive comparison between the averages across trials of the two conditions was not significant. Together, this suggests that the difference between early and late conditions is relatively small.

**Figure 2:**
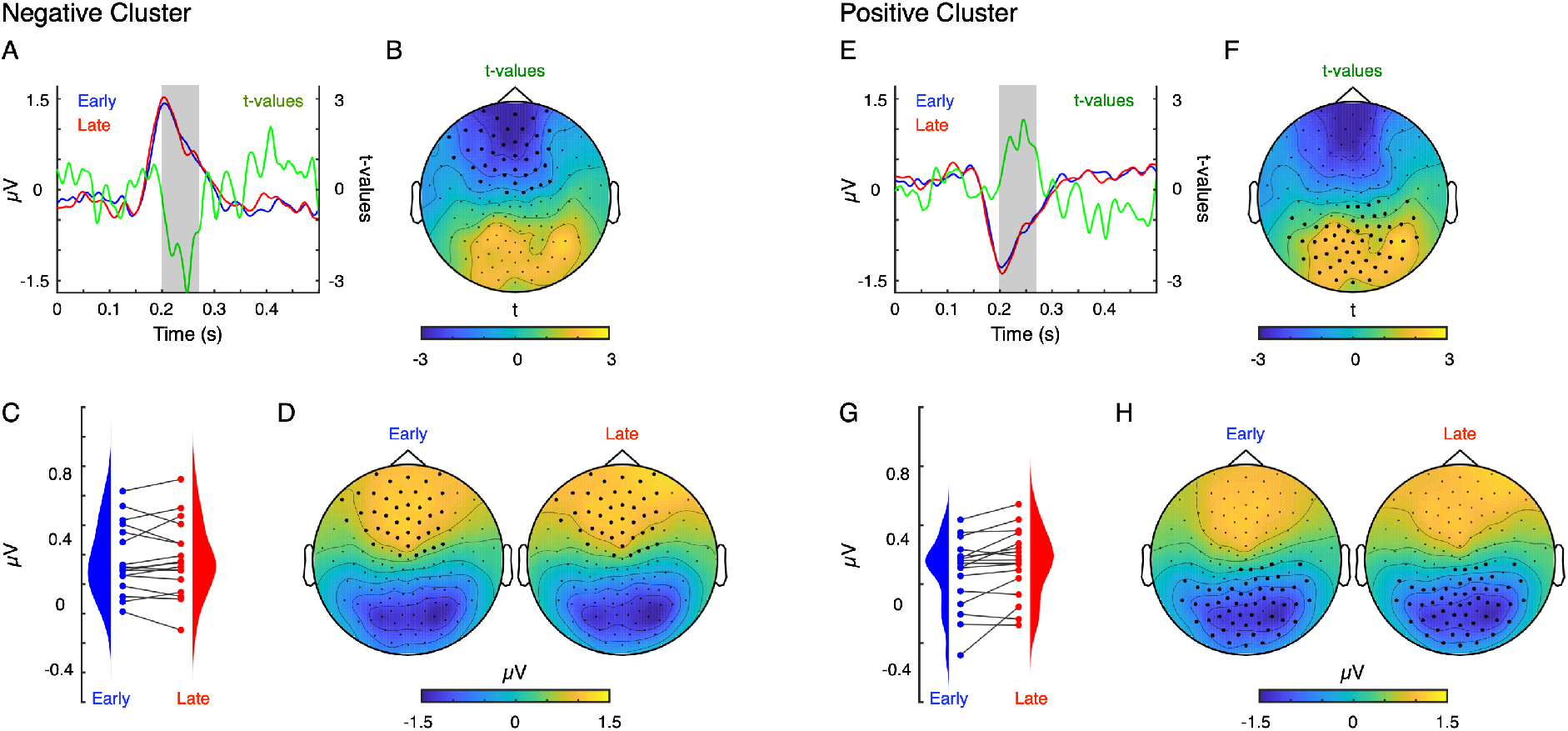
Differences between early and late flash-lag trials are reflected in early evoked responses. (A and D) Early (blue) and late (red) flash-lag trials evoked a positive peak around 200 ms after flash onset. In the comparison of the beta values from the linear model against zero across participants (green line), an early interval reflected the factor Time. (B) The early interval (200 – 276 ms) comprised a negative cluster across frontal electrodes. (C) Even though the beta values derived from single trials differed significantly from zero, the ERP amplitudes averaged across trials did not differ between early and late trials. (E-H) Same as A-D, but for the positive cluster across occipital electrodes in the same interval.

For the factor Illusion, the cluster-based permutation test revealed a late interval between 368 and 452 ms during which the beta values differed significantly from zero (Figure 3). This suggests significant single-trial ERPs differences between *illusion* and *no illusion* trials. Specifically, a cluster of negative beta values was found across central electrodes (cluster-p = 0.02 +/− 0.0087). Averaged across the negative cluster, trials in which the participants perceived the FLI were associated with a more positive ERP amplitude (Figure 3A blue,0.3803 +/− 0.2055) than trials in which participants did not perceive the illusion (Figure 3A red, 0.2339 +/− 0.2410). This difference was statistically significant (t(16) = 4.8825, CI = [0.0828 0.2100], p < 0.001, BF10 = 181.3138). Across participants the correlation between the ERP amplitude averaged over the cluster and the illusion rate was not significant (r(15) = 0.3333, p = 0.1910, BF10 = 0.4311).

**Figure 3:**
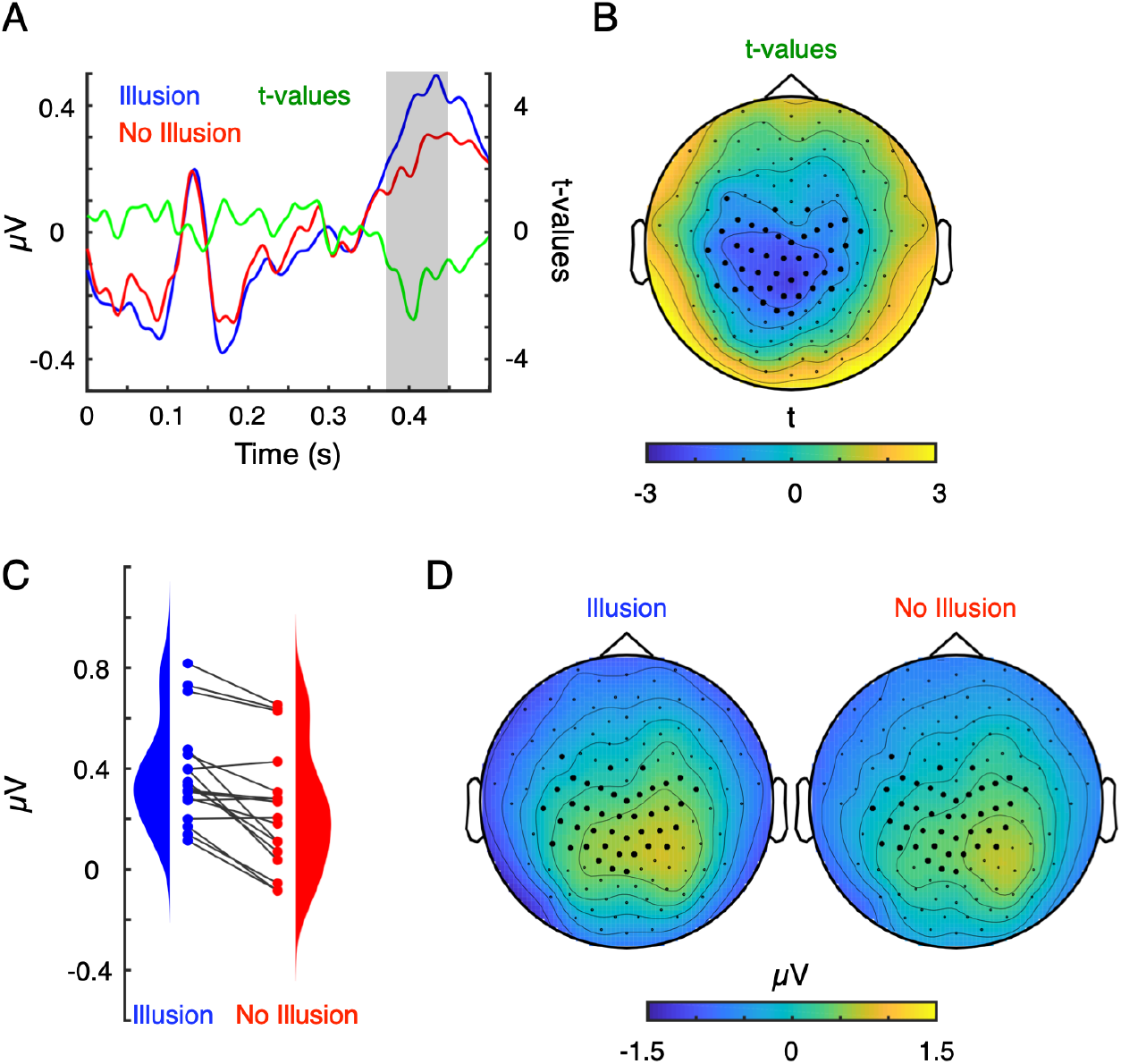
Differences between illusion and no illusion flash-lag trials are reflected in a late evoked response. (A and D) Illusion (blue) and no illusion (red) flash-lag trials evoked a positive peak around 400 ms after flash onset. In the comparison of the beta values from the linear model against zero across participants (green line), a late interval reflected the factor Illusion. (B) The late interval (368 – 452 ms) comprised a negative cluster across central electrodes. (C) The beta values derived from single trials differed significantly from zero, and the ERP amplitudes averaged across trials were more positive for illusion compared to no illusion trials.

The exploratory 2×2 repeated-measures ANOVA with the factors Time (early vs. late) and Illusion (*flash-lag* vs. *no flash-lag*) did not reveal any effects. Visual inspection of the uncorrected F and BF10 values indicates short-lived effects, and overall little support for the H1 outside the previously described effects (Supplementary Material).

### 3.3 Source Analyses

For the factor Time, the comparison between source space activity for *early* and *late* trials in the 200 to 276 ms interval after stimulus onset did not reveal any significant differences, which is in line with the ERPs analysis (averaged across trials) between conditions at the sensor level. For the factor Illusion the analysis of source space activity revealed an enhanced activity for *illusion* vs. *no illusion* trials in the 368 ms to 452 ms interval (cluster-p = 0.0313 +/− 0.0034; Figure 4). Comparison to the AAL atlas indicated the left inferior occipital gyrus as the primary source of the peak difference. Averaged across the nodes of the cluster, trials in which the participants perceived the FLI (1.6783 +/− 1.8015) were associated with stronger source space signal change from baseline than trials in which participants did not perceive the illusion (0.5888 +/− 0.8211). This difference was statistically significant (t(16) = 3.6991, CI = [0.4651 1.7139], p = 0.0019, BF10 = 21.3389). Across participants there was no significant correlation between the ERP amplitudes averaged over the cluster and the illusion rates (r(15) = 0.1961, p = 0.4492, BF10 = 0.2442).

**Figure 4:**
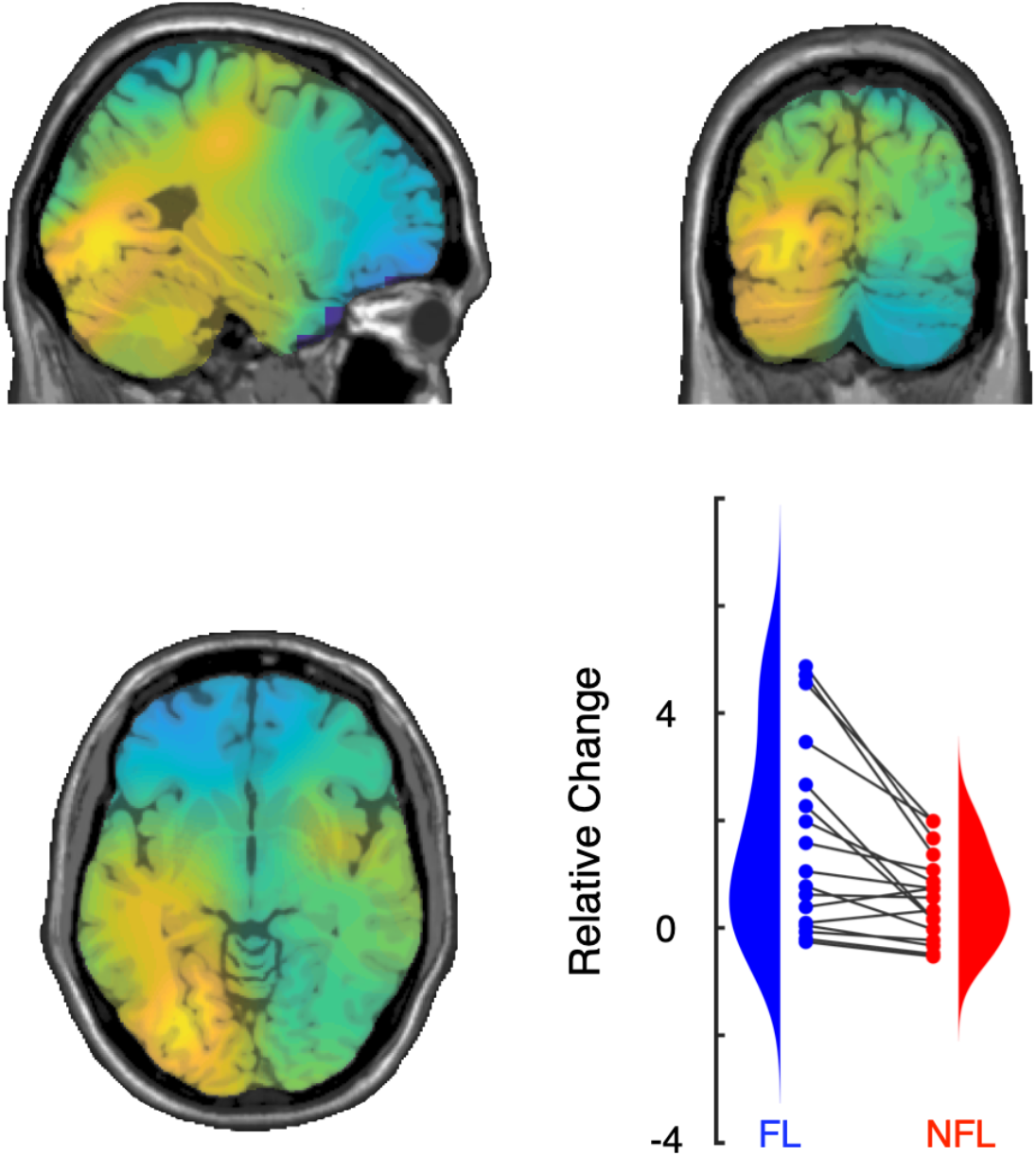
Source analysis for the factor Illusion revealed an involvement of the left fusiform gyrus. The comparison of source-space activity indicated an occipital cluster where trials in which participants perceived the flash-lag illusion evoked a stronger signal change from baseline than trials in which they did not perceive the illusion.

## 4 Discussion

In our study, we used ERPs to clarify the temporal and spatial neural correlates of the FLI. Some theories predict earlier, stimulus-dependent temporal processing, and others later stimulus-independent temporal processing. We conducted this analysis on a large pool of participants, from which we selected those with an approximately bistable perception of the illusion. A late positive ERP component (368-452ms after the flash onset), localized to the inferior occipito-temporal cortex differentiated between illusion and no illusion perception trials. Illusion trials evoked larger evoked cortical responses than trials in which no illusion occurred. Moreover, in our study, FLI perception was higher in late vs. early trials, with corresponding differences in an earlier ERP component (200-276ms).

We found evidence for the influence of late modulation in FLI, which indicates that late processing in the inferior occipital cortex is an important neural origin where perception in the FLI is reconstructed. Thus, later ERP components (>300ms, e.g. P300), which are typically associated with higher-order cognitive (Huang et al., 2015) or post-perceptual processes (Schröder et al., 2021) seem to be critical for the difference between illusion and no illusion perception, at least in our paradigm. This is consistent with a more global processing of visual information in the FLI, temporally decoupled from the stimulus sequence (Shimojo, 2014; Sergent, 2018). One possible interpretation of these results is that attention can modulate the strength of the FLI. For example, the FLI is increased when attentional resources are reduced, such as in a dual-task condition (Sarich et al., 2007) or spatial attention (Baldo et al., 2002). However, top-down attention typically produces enlarged ERPs components in primary sensory regions (Keil et al., 2017; Lange et al., 2008), which is difficult to reconcile with our results, in which larger ERP components are associated with FL rather than NFL. In summary, our findings show that late ERP components differentiate between illusion and no illusion perceptions, which supports the postdiction view on the FLI.

Our results are less congruent with theories that postulate early visual processing as critical to the FLI, such as the motion extrapolation (Hogendoorn, 2020) and reentrant based non-linear motion integration (Arstila, 2015). Visual inspection suggests small differences in early components, and Bayes Factors argues against the interpretation of a meaningful difference in early components < 200ms. However, it is also possible that there are important early effects that are too subtle to detect with our sample size. On the face of it, this is difficult to reconcile with the empirical findings that the FLI can be disrupted at approximately 200ms post flash by TMS stimulation of MT+ (Maus et al., 2013). However, this may mean that early activity is necessary for later processing of the visual stimuli, but not sufficient for the perception of the illusion, or that early activity is correlated with later processing rather than being causally necessary (Sergent, 2018). In support of this, we observe that there were differences between the early and late onset flashes at 200 ms to 276ms, which suggests that processing at this time relates to the registration of differences between the physical properties of the stimuli, rather than the FLI itself. Thus, interference with this physical coregistration of stimuli could disrupt upstream perception of the FLI. In all, our results do not appear to reflect the early stimulus-locked processing predicted by motion extrapolation or non-linear motion integration. The FLI dependent on later neural processing steps, decoupled from the temporal order of stimulus presentation.

Information regarding the cortical areas involved in the reconstruction of visual perception in the FLI come from our source analysis. This analysis highlighted important regions in FLI processing in approximately left fusiform gyrus. Activity in this region, including later ERP activity has been associated with a heterogenous group of spatial and temporal integrative phenomena. Previously, Meyer & Olson, (2011) found that inferotemporal cells in monkeys were sensitive to violations of statistical regularities in sequentially presented stimuli. Sequence representation has been generalized to encompass various linguistic phenomena (Dehaene et al., 2015). In a broader view, inferior temporal areas have been implicated as a part of a subcortical-extrastriate-fronto-parietal network necessary for the conscious perception of bistable stimuli (Bisenius et al., 2015). Integrating FLI with these models of sensory integration in this region could provide a new way of understanding the FLI phenomenon.

The study has some limitations. Our sample was restricted to those remaining participants with a more balanced number of illusion and no illusion trials, so it is possible that the current findings are not generalizable to the other participants. Future studies could adapt the flash onset to individual perception thresholds to ensure a larger sample size (as in e.g. Chakravarthi & VanRullen, 2012). Connected with this, the ERPs were based on a relatively small number of trials per person. It may also be that these factors contributed to the absence of ealier effects in the present study. Nevertheless, the sensitivity of the single-trial linear modelling combined with robust clustering statistics provide reason for confidence in the significant later ERP components. Our source analysis with EEG and non-individualized MRI templates is limited in spatial resolution, and future studies involving e.g. fMRI or MEG, would be necessary to strengthen these findings. Our primary stimuli went left to right, which is congruent with reading stimuli, this may in part explain the activation in the fusiform gyrus. Future studies could build on these findings by ERPs with different types of flash lag variants to test the generalizability of these findings. Furthermore, an attention manipulation could help unconfound potential effects (Koch & Tsuchiya, 2012; Koivisto, Kainulainen, & Revonsuo, 2009; Moran et al., in press). Overall, we believe that these limitations do not substantially affected our main finding that late evoked potentials reflect perception-related processing in the FLI. Nevertheless, they should be considered in further research studies examining the neural signatures of the FLI.

### 4.1 Conclusion

Our study shows for the first time the time-course of neural activity of the FLI, differentiating illusion from non-illusion trials. Although the different theories posited to explain FLI likely all have some purchase on the truth of this complex phenomenon, our results argue for a greater focus on later postdictive processing, decoupled from the order of stimulus presentation.

## Acknowledgments

This research was funded by a grant to DS from the German Research Foundation (SE1859/4-1). We thank Marianne Bröker, Alex Masurovsky, Teresa Ramme, Lisa Renziehausen, and Joseph Wooldridge for their assistance gathering the data.

## 6 Supplementary Material

The exploratory 2×2 repeated-measures ANOVA with the factors Time (*early* vs. *late*) and Illusion (*flash-lag* vs. *no flash-lag*) did not reveal any effects following FDR correction for multiple comparisons. Visual inspection of the uncorrected F and BF10 values indicates short-lived effects, and overall little support for the H1outside the previously described effects.

**Supplementary Figure:**
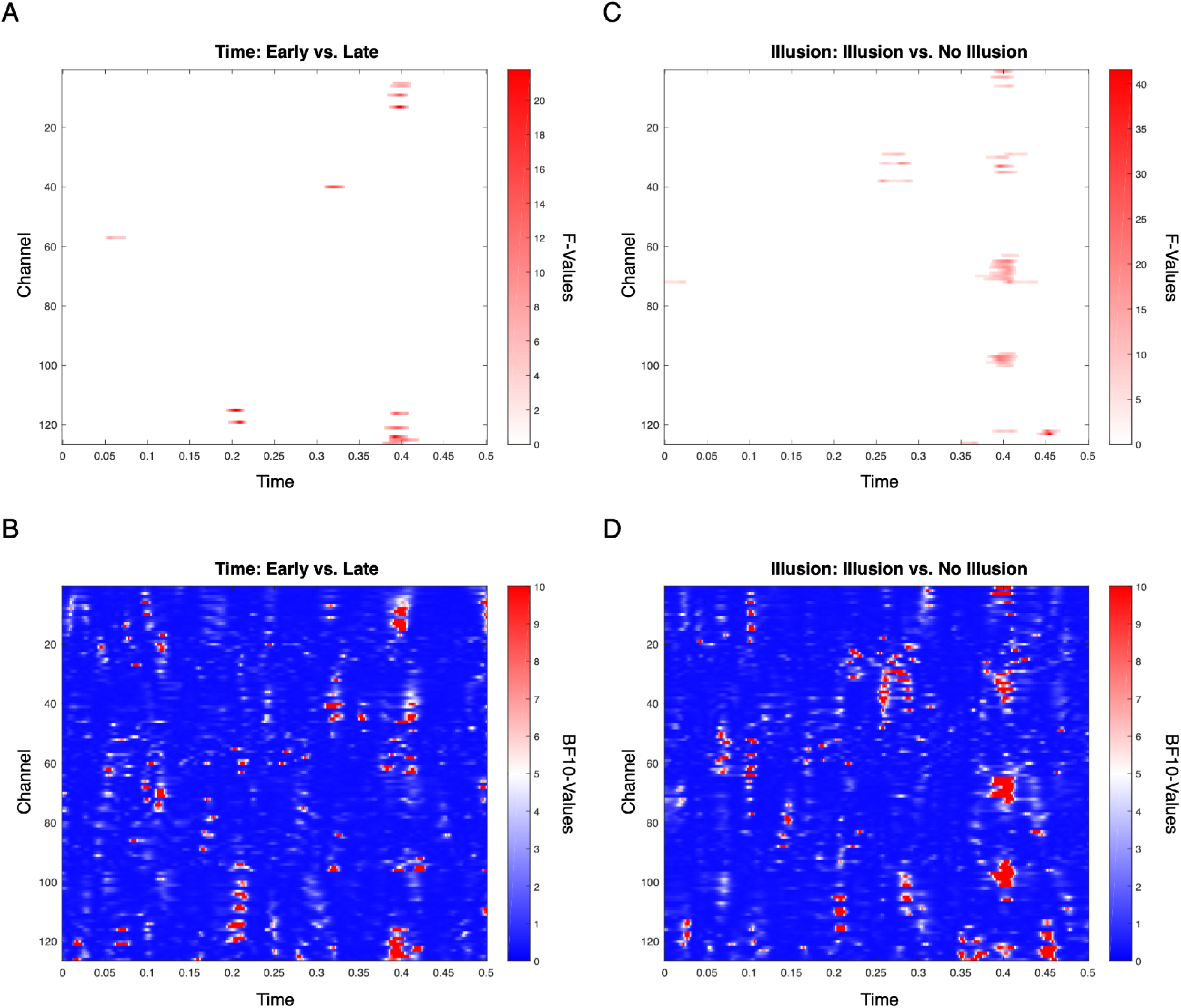
The exploratory 2×2 ANOVA did not reveal any effects outside the two intervals reported in the linear model analysis. (A) For the factor Time, F-values masked by the uncorrected threshold of p < 0.05 only reveal scattered peaks. (B) Similarly, BF10-values are low, indicating little support for the H1. (C and D) Same as A and B but for the factor Illusion.

